# Unexpected Long Range Actions of AAV8

**DOI:** 10.64898/2026.11.20.767967

**Authors:** Jessica M. Lahr, Julie Ly, Elif Erdem, Michael R. Williams

## Abstract

Adeno-associated virus vectors (AAVs) have broad utility in neuroscience. A variety of features, such as titer, capsid, enhancers and promoters, coding and non-coding elements, can be utilized to pursue diverse experimental ends. The net effects of titer and capsid have been used experimentally to pursue retrograde and anterograde studies, such as through the monosynaptic spread of high-titer AAV1. The literature on AAV8 is conflicting, with evidence for and against its long-range action outside of the infected starter population. Using the mouse corticopontocerebellar tract, we report that AAV8, when used at high titers, can be associated with long range recombination outside of the starter population. This phenomenon depended upon the viral titer and was influenced by the presence of other experimental injections. Surprisingly, separate injections of high-titer AAV8 recombinase and recombinase-reporter AAV8s gave rise to recombination across long distances in populations not known to be monosynaptically connected. Given the possibility of long range and emergent effects, we suggest that investigators empirically determine titer, capsid, and injection designs that reduce false positives and/or indirect effects.

## Introduction

Adeno-associated virus vectors (AAVs) are widely used for neural circuit mapping and manipulation. AAV serotypes vary in transduction properties, therefore capsid and viral titer are important experimental variables [1, 2]. Some AAV serotypes such as 1, 6, and 9, spread beyond the starter population in retrograde and/or anterograde directions [2-4]. The ability for AAV8 to spread transneuronally remains unclear; there are reports both in support of and against such spread [2, 5]. An increased scrutiny of experimental parameters versus AAV performance is important to prevent misinterpretations and to highlight useful or challenging properties of varying serotypes. Here, we sought to determine if AAV8 can spread long-range in the adult mouse brain using the corticopontocerebellar tract.

## Results

### AAV8 transneuronal spread assayed by Cre-reporter AAV8

An AAV8-Cre-dTomato (2.10E13 GC/mL) was injected into left basal pons (BPN), and an AAV8 nuclear Cre-Reporter (2.50E13) was injected into the right cerebellum of adult C57BL/6J mice. The AAV Cre-reporter infected cell nuclei are initially labeled with mCherry fluorescent protein and are converted to Green Fluorescent Protein (GFP) expression after Cre-mediated recombination; this reporter recombination was used to determine AAV8 spread from the BPN. Due to the per-cell multiplicity of episomal AAV genomes, only fully Green (vs Red *and* Green) labeled cells were considered recombined. Fluorescent protein expression in fixed tissues was examined by microscopy. Cerebellar Cre-reporter recombination was prevalent when AAV8-Cre was injected into the BPN (Figure 1A). As expected, basal Cre-reporter recombination in the cerebellum was sparse when saline was injected into the BPN (Figure 1B) and was absent when AAV8-Cre was injected into the BPN but saline was injected into the cerebellum (Figure 1C). The presence of cerebellar Cre-reporter recombination in this experiment (1A) suggests that AAV8 could mediate recombination outside of the pontine starter population. We did not observe robust Cre reporter GFP expression in the BPN, suggesting that there is not significant retrograde action of AAV8. Cerebellar Cre-reporter recombination was quantified as the fraction of reporter-expressing nuclei that expressed EGFP but not mCherry fluorescence. When AAV8-Cre was injected into the BPN, 20.0% (21/105), 11.6% (10/86) and 9.0% (10/111) of reporter-expressing nuclei were fully recombined (mean 13.6%, SD 5.7; n = 3 mice; Figure 1A). When saline was injected into the BPN, no fully recombined nucleus was detected in any animal (0/84, 0/80, 0/77; n = 3 mice; Figure 1B), placing the upper bound of the Cre-independent complete recombination rate at 1.52% (pooled 0/241 nuclei, 95% Clopper-Pearson interval). The two conditions were completely separated, with every AAV8-Cre animal exceeding every saline animal and differed significantly by a beta-binomial model accounting for between-animal variability (likelihood ratio test, parametric bootstrap p = 0.0082). Partially recombined nuclei, which expressed EGFP while retaining mCherry, occurred at comparable rates in the two conditions (12.9% vs 22.6%; p = 0.37), indicating that Cre-independent reporter activity does not differ between groups and thus cannot account for the fully recombined population. The density of reporter-expressing nuclei was similar across all six animals (1872-2698 nuclei per mm2), indicating that this quantification is not confounded by differences in reporter transduction.

**Figure 1.**
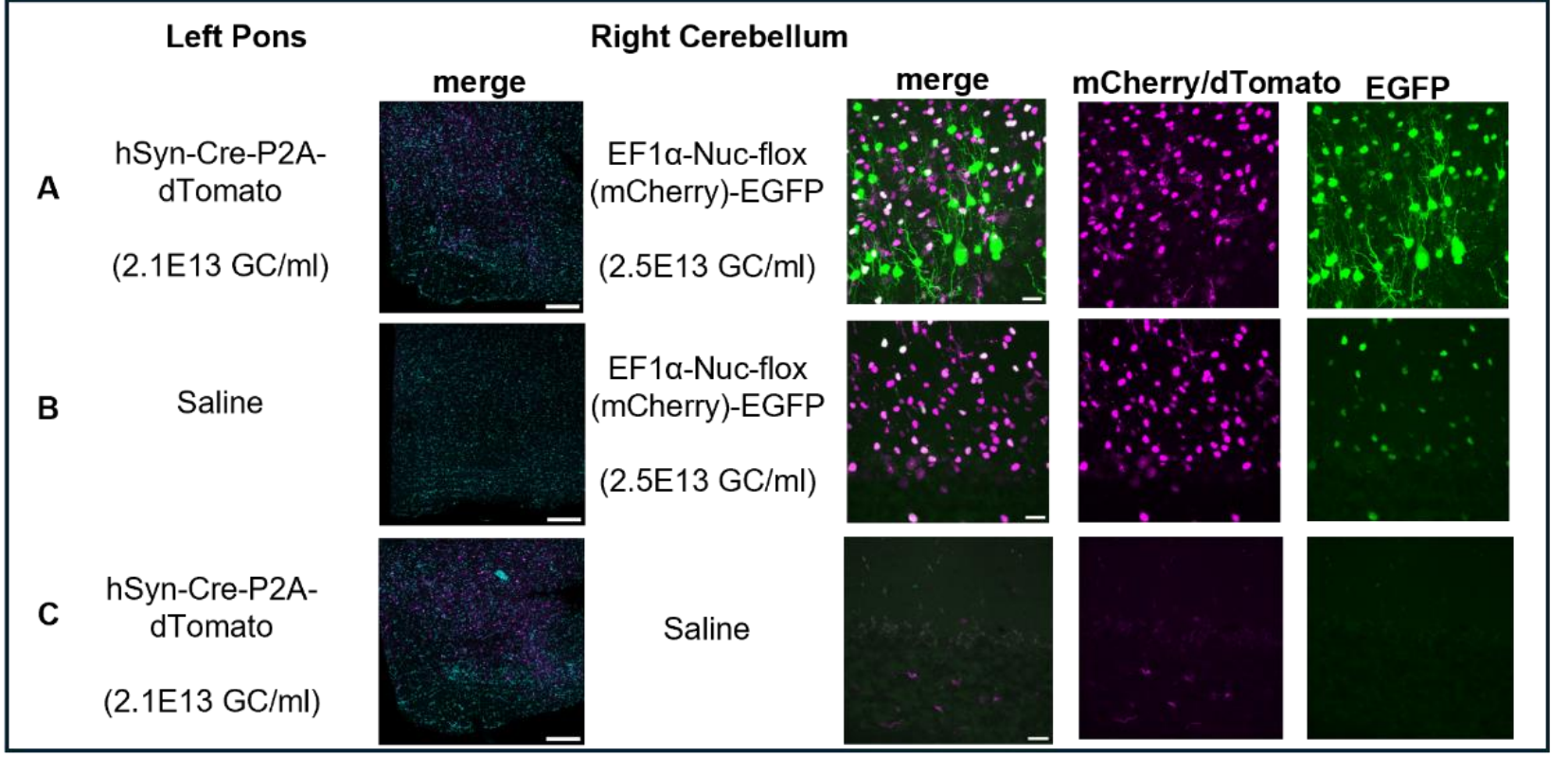
AAV8 anterograde spread from pons to contralateral cerebellum assessed by AAV8 (mCherry to EGFP) Cre-mediated recombination. Three adult mice were injected with each A) AAV8-Cre in the left pons and Cre-Reporter AAV8 in the cerebellum, B) saline in the left pons and Cre-Reporter AAV8 in the cerebellum, or C) AAV8-Cre in the left pons and saline in cerebellum. Pons merged DAPI and dTomato tiled images at 20x, 0.5x zoom, scale bars: 200μm. Cerebellum mCherry/dTomato and/or EGFP 20μm max projection confocal images at 63x magnification, 0.5x zoom, scale bars: 20μm.

### AAV8 transneuronal spread dependent on viral titer

The long-range action of some serotypes of AAVs is influenced by titer, such as the anterograde spread of AAV1. To determine if the observed recombination of AAV8 outside of the starter population was influenced by titer, differing titers of AAV8-Cre-dTomato were injected into the BPN of adult C57BL/6J mice (2.10E13, 2.10E12, 2.10E11), while the cerebellar injection of AAV8 nuclear Cre-reporter titer remained constant (2.50E13). The highest level of AAV8 spread (cerebellar reporter recombination) was observed at the E13 titer (Figure 2A), minimal spread at E12 (Figure 2B), and no spread at E11 (Figure 2C). These results indicate that the apparent anterograde spread of AAV8 mediated recombination is dependent on titer. Fully recombined nuclei were detected only at the highest titer: 20.0%, 11.6% and 9.0% of reporter-expressing nuclei at 2.10E13 (n = 3 mice; Figure 2A); 0%, 1.6% and 0% at 2.10E12, representing 2 of 295 nuclei in one of three mice (Figure 2B); and 0% in every animal at 2.10E11 (0 of 282 nuclei; Figure 2C). Recombination increased significantly with injected titer (beta-binomial model with log10 titer as a continuous covariate, slope 3.24 logit per log10 GC/mL, likelihood ratio test, parametric bootstrap p = 0.0022; Jonckheere-Terpstra trend test across saline and all three titers, p = 0.0014). Neither of the lower titers was individually distinguishable from saline (2.10E12, 2/295 vs 0/241 nuclei, Fisher exact p = 0.50; 2.10E11, 0/282 vs 0/241 nuclei, identical).

**Figure 2.**
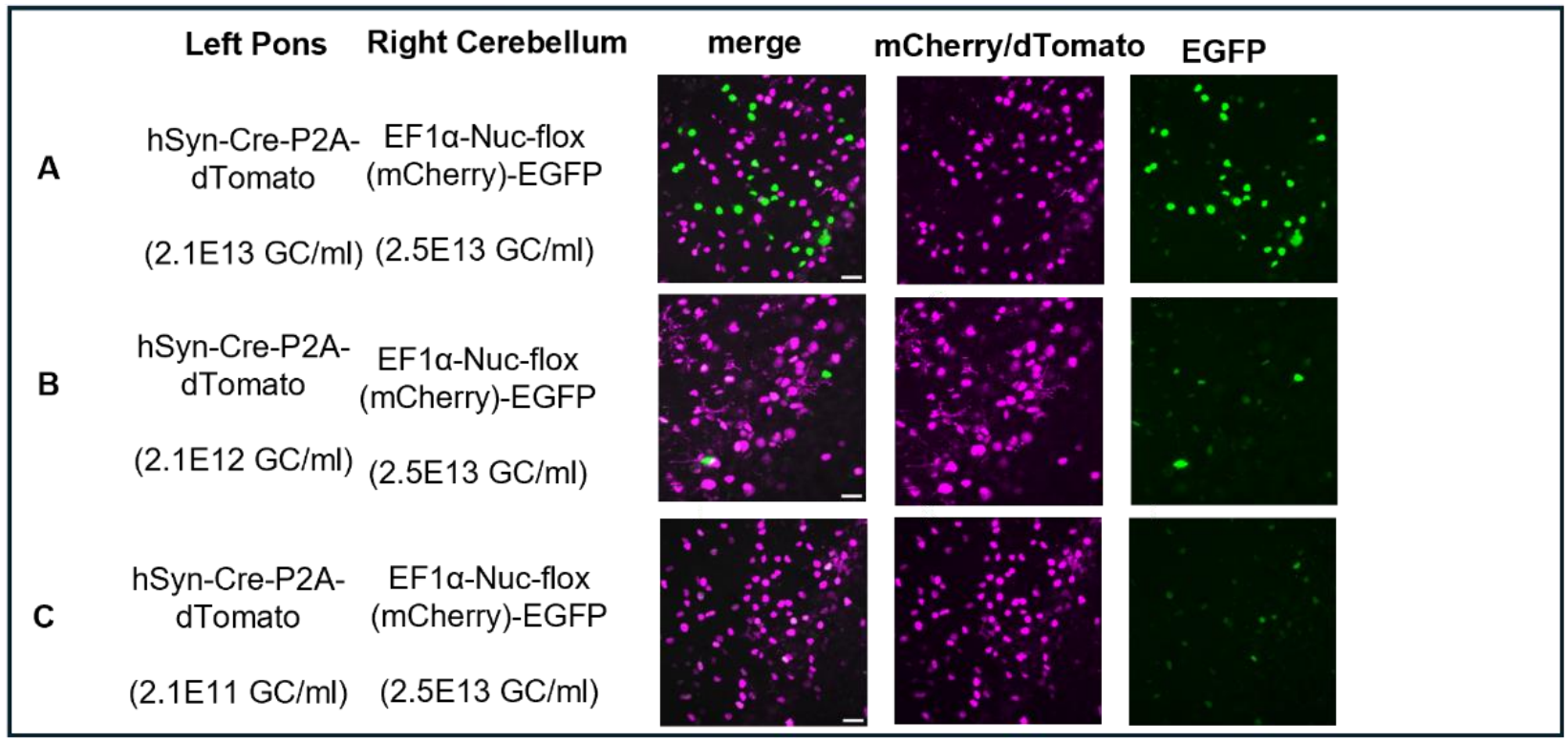
Titer dependence of AAV8 spread assessed by AAV8 (mCherry to EGFP) Cre-mediated recombination in contralateral cerebellum vs different titers injected into pons. Three adult mice were injected with each A) AAV8-Cre in the left pons at 2.1e13 GC/ml and Cre-reporter AAV8 in the cerebellum B) AAV8-Cre in the left pons at 2.1e12 GC/ml and Cre-reporter AAV8 in the cerebellum C) AAV8-Cre in the left pons at 2.1e11 GC/ml and Cre-reporter AAV8 in the cerebellum. Cerebellum mCherry/dTomato and/or EGFP 20μm max projection confocal images at 63x magnification, 0.5x zoom, scale bars: 20μm.

### Long range AAV8 action and secondary manipulations

We sought to determine if the apparent long-range action of high-titer AAV8 depended upon the reporter system. We repeated high-titer AAV8 Cre injection into the BPN of Ai14 (Cre-on TdTomato) mice. Even at 2.10E13 AAV8 Cre titers into the BPN, we did not observe robust TdTomato reporter allele induction in cerebellar cells (Figure 3A). However, with a second manipulation, whether saline, or Cre-reporter AAV8 injected into the cerebellum, the Ai14 reporter and/or viral reporter expression were then observed in cerebellum (Figure 3B,C). When saline (instead of AAV8 Cre) was injected into BPN, there was only the expected sparse low “background level” of recombination of cerebellar Cre reporters (Figure 3D). These results suggest that for high titer AAV8, secondary injections to, or AAV exposure in, secondary locations can facilitate recombination outside the starter population.

**Figure 3.**
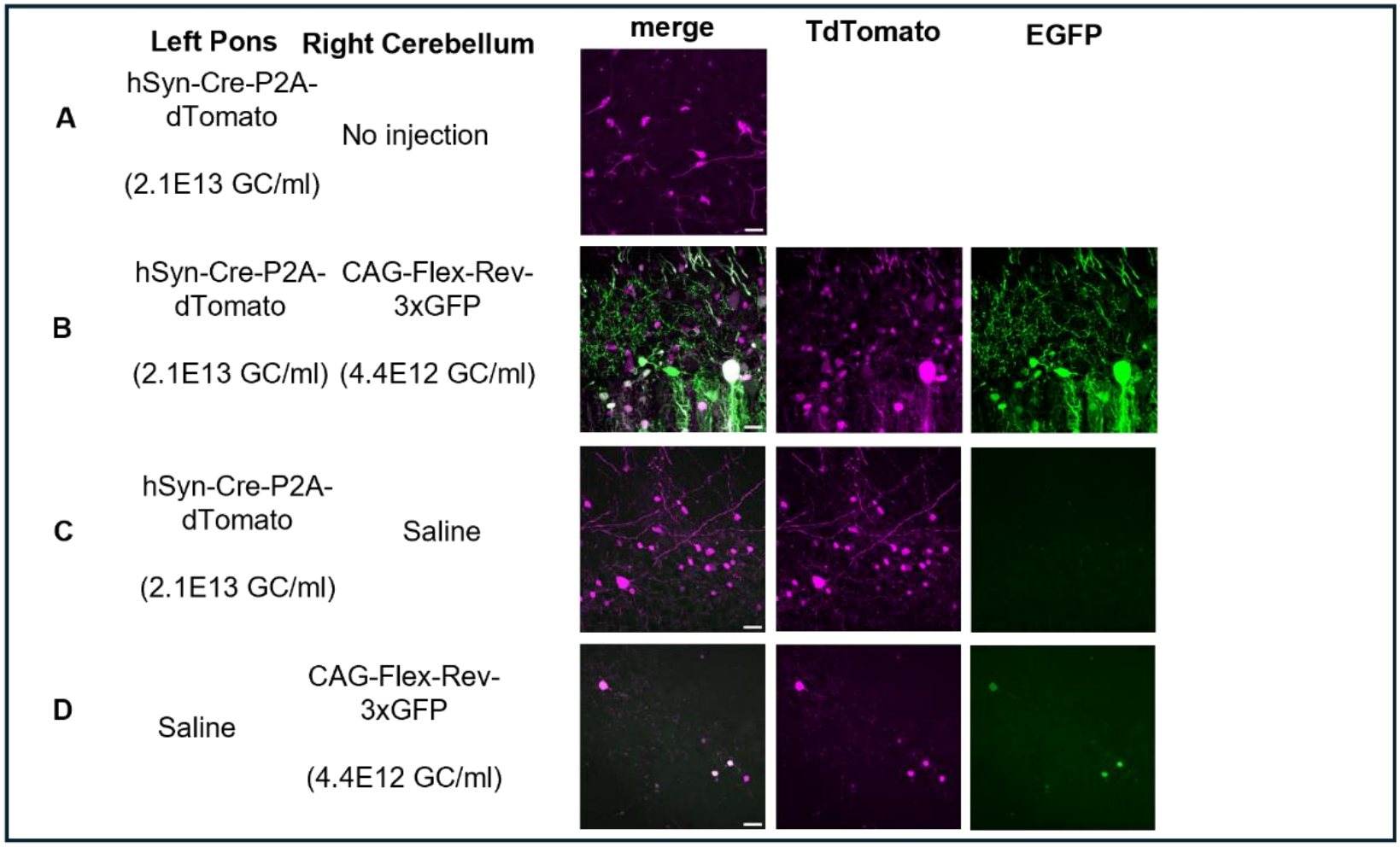
AAV8 spread from pons to contralateral cerebellum, assessed by Ai14 Cre-On TdTomato allele and/or AAV8 Cre-On EGFP recombination. Three adult Ai14 Cre-reporter mice were injected with each A) AAV8-Cre into the left pons with no injection in cerebellum, B) AAV8-Cre in the left pons and Cre-reporter AAV8 in cerebellum, C) AAV8-Cre in the left pons and saline in cerebellum, or D) saline in the left pons and AAV8-Cre reporter in cerebellum. Cerebellum mCherry/TdTomato and/or EGFP 20μm max projection confocal images at 63x magnification, 0.5x zoom, scale bars: 20μm.

### Long-range AAV8 spread across noncanonical neural circuit

The recombination of an AAV Cre reporter in the cerebellum following high-titer AAV8 Cre expression in the BPN suggested transneuronal anterograde spread of AAV8. However, such recombination was widespread (Figures 1A and 3B) and was not restricted to granule cells (monosynaptic anterograde targets of pontine mossy fibers). Therefore, other mechanisms may explain long-range recombination. We sought to determine if AAV8 could act over long-range from the V1 visual cortex to the cerebellum, since these regions are not known to be monosynaptically connected. AAV8-Cre (2.10E13) was injected into left V1 and Cre-reporter AAV8 was injected into right cerebellum. We observed reporter recombination in the cerebellum when the AAV8 Cre was injected into the BPN (Figure 4). Cerebellar Cre-reporter recombination following V1 injection was 19.8% (17/86), 8.1% (5/62) and 13.2% (9/68) of reporter-expressing nuclei (mean 13.7%, SD 5.9; n = 3 mice; Figure 4), significantly above the saline-injected control condition (beta-binomial likelihood ratio test, parametric bootstrap p = 0.0032; controls pooled 0/241 nuclei, upper 95% bound 1.52%). Recombination frequencies after V1 (Figure X) and BPN (Figure 1A) injection were indistinguishable (difference -0.6 percentage points, 95% bootstrap CI -8.5 to +7.1); at three mice per group this interval is wide, so these data establish that V1 injection produces recombination of the same order as BPN injection rather than that the two are equal. Cre-independent double-labeled nuclei did not differ between the two injection sites (p = 0.27). These results suggest that injections of high titer AAV8 Cre expressing and Cre reporting constructs may lead to recombination across long distances in neuronal populations not known to be monosynaptically coupled.

**Figure 4.**
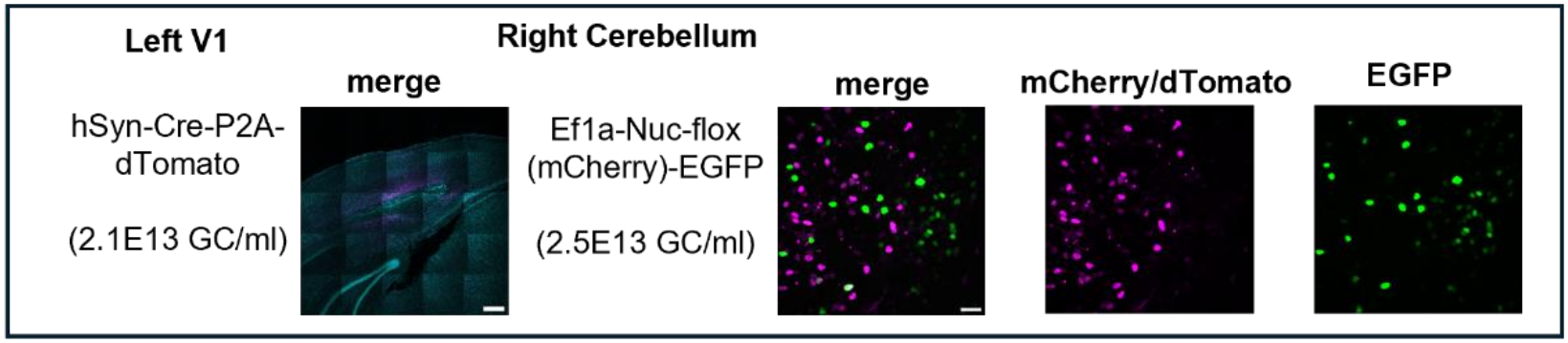
AAV8 spread from V1 to contralateral cerebellum, assessed by AAV8 (mCherry to EGFP) Cre-reporter recombination. Three adult mice were injected with AAV8-Cre in the left primary visual cortex (V1) and AAV8 Cre-reporter in the cerebellum. V1 injection site with merged DAPI and dTomato tiled images at 20x, 0.5x zoom, scale bar: 300μm. Cerebellum merged and separate mCherry/dTomato and EGFP 20μm max projection confocal images at 63x magnification, 0.5x zoom, scale bar: 20μm.

## Discussion

We find that AAV8 appears to spread to, and/or to induce recombination of reporter constructs at, cell populations distal from its injection location. The local vs long range action of AAV8 may depend upon the reporter system. Here, as in many studies, combinations of AAV8 recombinase and recombinase reporter injection sites were used, whereas some other previous studies variably used a single injection site and/or examined reporter alleles or non-AAV vectors [1, 2, 5]. Methodological differences may thus be relevant to the differing effects observed here versus other studies. Indeed, following injection of AAV8-Cre in an Ai14 reporter mouse with a single viral injection into the pons, we did not observe robust reporter recombination in the cerebellum (beyond starter pontine mossy fibers) and therefore, our results are not incompatible with former reports that observed no long-range action of AAV8. We did, however, observe apparent AAV8 spread from the pons when saline was injected into the cerebellum, and yet more apparent spread when a second AAV8 was injected into the cerebellum. Thus, the combination of high-titers *and* secondary perturbations may facilitate long-range AAV8 action outside of the starter population. It has been reported that cells undergoing stress may have increased endocytosis, so it is possible that the secondary injection in the receiving population enhances cellular endocytosis or nuclear import of AAV8-Cre, facilitating recombination [6, 7].

Unexpectedly, we found long-range AAV8 action between the V1 and cerebellum, which are not known to be monosynaptically connected [8]. While it is possible that there was leakage of AAV8 Cre virus into the cerebrospinal fluid, it would be surprising if such a small amount of vector would cause detectable recombination at the levels and distances observed, and it seems highly unlikely that such action would be coincidentally limited to the site of the secondary injection. Since the long-range action of high titer AAV8 that we report here is not limited to monosynaptically coupled anterograde populations, it is possible that AAV8 spreads in a polysynaptic anterograde manner, through non-synaptic transcellular trafficking, and/or through extracellular routes such as along fiber bundles, astrocytic networks, glymphatic or vascular tracts, or through other interstitial spaces [9]. Ultimately, more research is needed to determine the mechanisms for long-range action of AAV8 and to determine if AAV8 can exert long-range actions in other brain areas or species.

Our findings suggest that unexpected or spurious patterns of recombination could arise when using AAV8 experimentally. We do not, however, suggest that the utility of AAV8 is limited. Rather, we suggest that investigators empirically determine AAV titer ranges and reporter system experimental designs that are robust to false positives. These controls, and others such as empty capsid and/or vehicle injections, may especially be important when interpreting experiments that include multiple injections and/or that rely upon sparse phenomena.

## Methods

### Animals

Experiments were approved by the Institutional Animal Care and Use Committees of Michigan State University, which is accredited by the Association for Assessment and Accreditation of Laboratory Animal Care. All mice were on a 12-hour dark/light cycle, with ad libitum access to food and water. Adult male and/or female mice were used in each experiment. Mouse strains used for experiments were C57BL/6J (RRID:IMSR_JAX:000664) and Ai14 (RRID:IMSR_JAX:007914).

### Viral vectors

AAV particles used for stereotaxic injections were either purchased from Addgene or from the BRAIN Initiative Viral Vector Core Facility at the University of North Carolina Neuroscience Center (RRID:SCR_023280). Viruses were stored as 5ul aliquots upon arrival at -80C and were thawed on ice prior to injections.

### Stereotaxic viral injections

Injections were performed largely as previously described [10]. Mice were anesthetized using inhaled isoflurane (2-3%) (SomnoSuite, Kent), with thermal regulation by a heating pad and pain management through topical lidocaine (0.5%) and intraperitoneal ketoprofen (3 mg/kg). Viral vector injections were performed using a digital stereotaxic device (Kopf). Small drill holes were localized in the skull and viral particles were delivered using 30-gauge Hamilton Syringes that were driven by a digital syringe pump (WPI). Separate syringes were used for Cre and Cre reporter vectors. Each virus was delivered at a flow rate of 50 nl/min; the total delivered volume of each viral injection was between 200 and 250 nl. Unless noted, viral titers were as provided in Table 1. The needle remained in place for 5 minutes following viral infusion before being slowly withdrawn. Post-operative monitoring included intraperitoneal ketoprofen (3 mg/kg) 24- and 48-hours following surgery.

**Table 1.** AAV8 Virus name.

|  | **Titer (GC/ml)** |
|---|---|
| pAAV-hSyn-Cre-P2A-dTomato (Addgene, 107738-AAV8) | 2.10E13 |
| pOTTC1032 - pAAV EF1a Nuc-flox(mcherry)-EGFP (Addgene, 112677) | 2.50E13 |
| pAAV-CAG-Flex-Rev-3xGFP (Neurotools) | 4.40E12 |

Coordinates for injections into the Pons were A/P, -4.00 mm; M/L, -0.5 mm; D/V, 5.40-5.10 mm. Cerebellar coordinates were A/P, -6.35 mm; M/L, +1.6 mm; D/V, 1.5-0.75 mm. Primary visual cortex coordinates were A/P, -3.5 mm; M/L, -2.5 mm; D/V, 0.8-0.5 mm.

### Histology and Imaging

At 3-4 weeks post injection, avertin-anesthetized mice underwent transcardial perfusions and post-fixation with 4% paraformaldehyde as previously described [10]. Brains were sliced at 50 um using a vibratome (Leica, VT-1200S). Slices of brain regions of interest were mounted onto glass slides; a subset underwent DAPI staining. Representative regions of viral expression were imaged by confocal microscope (Zeiss LSM 800). In control experiments where saline was used, comparable anatomical regions were imaged. For each animal, 20x single plane and a 63x (20 um z-stack) images were collected at the anterograde (reporter) injection site. Tiled 20x single plane images were collected for the starter injection sites. For select conditions, a 20x single plane image was taken in the retrograde location of the circuit of interest. Each experimental condition included at least three animal replicates. Image processing was performed using FIJI/ImageJ

### Image quantification and statistical analysis

Recombination was quantified from the 63x maximum-intensity projections, acquired with constant laser power and detector gain across all animals. Channels were background-corrected by subtraction of a Gaussian-blurred estimate (15 um sigma). Nuclei were segmented from the union of the mCherry and EGFP channels by Li thresholding followed by watershed separation. The denominator throughout is reporter-expressing nuclei rather than total cells. A nucleus was scored as fully recombined if background-subtracted EGFP exceeded 57.3 arbitrary units with zero residual mCherry, and as partially recombined if it exceeded threshold while retaining mCherry; the EGFP threshold was set from the bimodal intensity gap in the first image analyzed and applied unchanged thereafter. One field (202.8 um square, 72-129 nuclei) was analyzed per animal, and the animal was the unit of analysis throughout. Per-animal counts were compared using a beta-binomial generalized linear model with significance from a likelihood ratio test calibrated by parametric bootstrap (4000 simulations). Titer dependence was tested by adding log10 titer as a continuous covariate and, independently, by the Jonckheere-Terpstra trend test. Proportions are reported with 95% Clopper-Pearson intervals. Analyses were performed in Python 3.13 using NumPy, SciPy, scikit-image and pandas. The V1 experiment was quantified by the same pipeline.

## References

1. Witteveen I, Balmer T. Comparative Analysis of Six Adeno-Associated Viral Vector Serotypes in Mouse Inferior Colliculus and Cerebellum. eNeuro. 2024 Nov 4;11(11):ENEURO.0391-24.2024. doi: 10.1523/ENEURO.0391-24.2024. PMID: 39467650; PMCID: PMC11576142.

2. Zingg B, Chou XL, Zhang ZG, Mesik L, Liang F, Tao HW, Zhang LI. AAV-Mediated Anterograde Transsynaptic Tagging: Mapping Corticocollicular Input-Defined Neural Pathways for Defense Behaviors. Neuron. 2017 Jan 4;93(1):33–47. doi: 10.1016/j.neuron.2016.11.045. Epub 2016 Dec 15. PMID: 27989459; PMCID: PMC5538794.

3. Salegio EA, Samaranch L, Kells AP, Mittermeyer G, San Sebastian W, Zhou S, Beyer J, Forsayeth J, Bankiewicz KS. Axonal transport of adeno-associated viral vectors is serotype-dependent. Gene Ther. 2013 Mar;20(3):348–52. doi: 10.1038/gt.2012.27. Epub 2012 Mar 15. PMID: 22418061; PMCID: PMC3381869.

4. Löw K, Aebischer P, Schneider BL. Direct and retrograde transduction of nigral neurons with AAV6, 8, and 9 and intraneuronal persistence of viral particles. Hum Gene Ther. 2013 Jun;24(6):613–29. doi: 10.1089/hum.2012.174. PMID: 23600720; PMCID: PMC3689167.

5. Castle MJ, Gershenson ZT, Giles AR, Holzbaur EL, Wolfe JH. Adeno-associated virus serotypes 1, 8, and 9 share conserved mechanisms for anterograde and retrograde axonal transport. Hum Gene Ther. 2014 Aug;25(8):705–20. doi: 10.1089/hum.2013.189. Epub 2014 May 2. PMID: 24694006; PMCID: PMC4137353.

6. Paul D, Stern O, Vallis Y, Dhillon J, Buchanan A, McMahon H. Cell surface protein aggregation triggers endocytosis to maintain plasma membrane proteostasis. Nat Commun. 2023 Feb 28;14(1):947. doi: 10.1038/s41467-023-36496-y. PMID: 36854675; PMCID: PMC9974993.

7. Zengel J, Carette JE. Structural and cellular biology of adeno-associated virus attachment and entry. Adv Virus Res. 2020;106:39–84. doi: 10.1016/bs.aivir.2020.01.002. Epub 2020 Feb 13. PMID: 32327148.

8. Wiesendanger R, Wiesendanger M. The corticopontine system in the rat. II. The projection pattern. J Comp Neurol. 1982 Jul 1;208(3):227–38. doi: 10.1002/cne.902080303. PMID: 7119159.

9. Cooper ML, Selles MC, Cammer M, Redd C, Gildea HK, Sall J, Chiurri KE, Cheung P, Wheeler DG, Saab AS, Liddelow SA, Chao MV. Astrocytes connect specific brain regions through plastic networks. Nature. 2026 Jul;655(8121):183–191. doi: 10.1038/s41586-026-10426-6. Epub 2026 Apr 22. PMID: 42020738; PMCID: PMC13322983.

10. Zaman T, Vogt D, Prokop J, Alsabia QA, Simms G, Stafford A, Luikart BW, Williams MR. Kit Ligand and Kit receptor tyrosine kinase sustain synaptic inhibition of Purkinje cells. Elife. 2024 Mar 27;12:e89792. doi: 10.7554/eLife.89792. PMID: 38536959. PMCID: PMC10972566.

